# Differential expression levels of Sox9 in early neocortical radial glial cells regulate the decision between stem cell maintenance and differentiation

**DOI:** 10.1101/2020.12.09.417931

**Authors:** Jaime Fabra-Beser, Jessica Alves Medeiros de Araujo, Diego Marques-Coelho, Loyal A. Goff, Ulrich Müller, Cristina Gil-Sanz

## Abstract

Radial glial progenitor cells (RGCs) in the dorsal forebrain directly or indirectly produce excitatory projection neurons and macroglia of the neocortex. Recent evidence shows that the pool of RGCs is more heterogeneous than originally thought and that progenitor subpopulations can generate particular neuronal cell types. Using single cell RNA sequencing, we have studied gene expression patterns of two subtypes of RGCs that differ in their neurogenic behavior. One progenitor type rapidly produces postmitotic neurons, whereas the second progenitor remains relatively quiescence before generating neurons. We have identified candidate genes that are differentially expressed between these RGCs progenitor subtypes, including the transcription factor Sox9. Using *in utero* electroporation, we demonstrate that elevated Sox9 expression in progenitors prevents RGC division and leads to the generation of upper-layer cortical neurons from these progenitors at later ages. Our data thus reveal molecular differences between cortical progenitors with different neurogenic behavior and indicates that Sox9 is critical for the maintenance of RGCs to regulate the generation of upper layer neurons.

**SIGNIFICANCE STATEMENT:** The existence of heterogeneity in the pool of RGCs and its relationship with the generation of cellular diversity in the cerebral cortex has been an interesting topic of debate for many years. Here we describe the existence of a subpopulation of RGCs with reduced neurogenic behavior at early embryonic ages presenting a particular molecular signature. This molecular signature consists of differential expression of some genes including the transcription factor Sox9, found to be a specific master regulator of this subpopulation of progenitor cells. Functional experiments perturbing Sox9 expression’s levels reveal its instructive role in the regulation of the neurogenic behavior of RGCs and its relationship with the generation of upper layer projection neurons at later ages.

## INTRODUCTION

The neocortex is evolutionarily the most recent addition to the brain and executes complex tasks including sensory integration and cognition. Neocortical development depends on precisely orchestrated mechanisms ranging from the generation of the neural cells in specialized germinal zones, to their migration to defined locations and the establishment of specific connections. During cortical development radial glia progenitor cells (RGCs) generate different subtypes of excitatory neurons that will ultimately form the layers of the cerebral cortex and make connections to distinct subcortical and cortical targets. The mature cortex consists of six principal layers that are generated following a temporal sequence. Neurons that populate deep layers of the cortex are generated first, followed by upper layer neurons and finally glia cells (Molyneaux et al., 2007).

Multiple progenitor subtypes with distinct morphologies and behaviors have been described in the neocortex. RGCs occupy the ventricular zone (VZ) and are connected to the ventricular and pial surfaces of the neocortex by apical and basal processes, respectively. Intermediate progenitor cells (IPCs) are basal progenitors grouped in the subventricular zone (SVZ) and are mostly generated by asymmetric divisions of RGCs (Miyata et al., 2004; Noctor et al., 2004). Other types of progenitor cells have been described more recently including short neural precursors (SNP) (Gal et al., 2006) and outer radial glial cells (oRGC) (Fietz et al., 2010; Hansen et al., 2010; Reillo et al., 2011). oRGCs are expanded in gyrencephalic animals and are thought to be important for the expansion of neocortical neurons in primates, especially those occupying the enlarged upper cortical layers (Lui et al., 2011; LaMonica et al., 2012).

Despite the advance of knowledge in the recent years in the identification of morphological differences among neocortical progenitors, the mechanisms that regulate neuronal progenitor diversification are not well understood. In fact, the extent of neocortical progenitor heterogeneity is still unclear and, in spite of the significant effort in the last 50 years to investigate cortical development, it is still unknown how the population of neocortical progenitors is able to generate the diversity of neocortical cells. The traditional hypothesis states that the fate of each neuronal subtype is determined by its birthdate, suggesting the existence of a common RGC progenitor that becomes progressively restricted with developmental time (McConnell and Kaznowski, 1991; Frantz and McConnell, 1996; Desai and McConnell, 2000; Shen et al., 2006; Gao et al., 2014). However, recent studies suggest that the pool of RGCs is more diverse, with some RGCs primarily generating certain types of cells including upper layer cortical neurons and B1 cells (Franco et al., 2012; Fuentealba et al., 2015; García-Moreno and Molnár, 2015; Gil-Sanz et al., 2015; Llorca et al., 2019). Nevertheless, the mechanisms that might lead to progenitor diversification are unclear, and molecular markers that would distinguish disparate populations of RGCs are largely unknown.

To identify molecular differences between RGC progenitors, we used an *in utero* electroporation approach to introduce plasmid constructs containing different promoters linked to reporter genes, based on the prediction that different promoters may be active in distinct neocortical RGC subtypes as described (García-Moreno and Molnár, 2015). Using this strategy, we identified a subpopulation of RGCs with reduced neurogenic behavior at early embryonic ages that eventually produces predominantly cortico-cortical projection neurons occupying upper neocortical cell layers. Using single cell RNA sequencing (scRNA seq), we characterized the molecular signature of these RGC progenitors and identified by transcriptional regulatory network analysis that they express *Sox9* at significantly higher levels compared to other RGCs. Notably, Sox9 is a transcription factor involved in stem cell maintenance (Scott et al., 2010; Liu et al., 2016) and associated with progenitor quiescence inside and outside the nervous system (Kadaja et al., 2014; Llorens-Bobadilla et al., 2015). Using functional perturbations, we demonstrate that Sox9 is critical to control self-renewal of a subset of RGCs to preserve them during early stages of corticogenesis for the production of upper layer neocortical neurons at later developmental stages.

## MATERIALS AND METHODS

### Mice

All experiments involving mice were conducted in accordance with the Spanish legislation as well as with the Directive 2010/63/EU of the European legislation and the United States of America animal welfare regulations and were approved by the ethical committee of the University of Valencia and the Conselleria de Agricultura, Desarrollo Rural, Emergencia Climática y Transición Ecológica of the Comunidad Valenciana and Institutional Animal Care and Use Committee (IACUC) of the USA.

C57BL/6 wild-type (WT) and Ai9 Cre reporter strains (Madisen et al, 2010) used for experimentation were originally obtained from the Jackson laboratory and are currently bred in the animal facilities of the University of Valencia, placed in the Burjassot campus. Both female and male mice were analysed in this study. Induction of recombination in *Ai9* mice electroporated with Cre inducible plasmids was achieved by intraperitoneal injection of pregnant dams with the faster-acting and shorter-lived tamoxifen metabolite 4-hydroxy-tamoxifentamoxifen (4-OHT) (Sigma) (1 mg/20 g of body weight, dissolved as described (Guenthner et al., 2013)).

### Expression Constructs

Hes5p-dsRed (Addgene plasmid #26868) was generated by Nicholas Gaiano (Mizutani et al., 2007). Hes5p-GFP was generated by replacing dsRed2 with eGFP in Hes5p-DsRed (Franco et al., 2012). pCAG-mRFP1 (Addgene plasmid #32600) was deposited by Anna-Katerina Hadjantonakis (Long et al., 2005). pCAG-mRFP;Hes5p-GFP was generated by subcloning pCAG-mRFP and a poly-adenylation signal into Hes5p-GFP. pCAG-CreERT2 (Addgene plasmid #14797) was deposited by Connie Cepko (Matsuda and Cepko, 2007). Hes5p-CreERT2 was generated by replacing pCAG with Hes5p. pCIG (Addgene plasmid #11159) was deposited by Connie Cepko (Matsuda and Cepko, 2004). CBFRE-GFP (Addgene plasmid #17705) was deposited by Nicholas Gaiano (Mizutani et al., 2007). TOP-GFP (Addgene plasmid #35489) was deposited by Ramesh Shivdasani (Horst et al., 2012). EMX2p-GFP was generated according to the publication of García-Moreno and Zoltán Molnár (2015). pCIG-S9FL was generated by cloning *Sox9* coding sequence into pCIG.

### *In Utero* Electroporation

*In utero* electroporation was performed as described (Gil-Sanz et al., 2013). Briefly timed pregnant mice were anesthetized with isoflurane. Once the pedal reflex was lost, analgesic solution was injected subcutaneously. The abdominal area was shaved, and a Caesarean section was performed to expose the uterine horns. 0.5-1 μL of endotoxin-free plasmid DNA solution (~1 μg/μL) was injected into one of the embryos’ lateral ventricles. For electroporation of embryonic day (E) 12.5 embryos, five pulses of 30 V were delivered, spaced by 950ms. After electroporation, the uterus was placed in the abdominal cavity, the abdominal wall and skin were sutured. Embryos were let to develop *in utero* for the indicated time or females were allowed to give birth and pups were sacrificed at the indicated age.

### *In Utero* Flashtag Injection

Carboxyfluorescein succinimidyl ester or FlashTag (FT) injections were performed at E14.5 as described (Govindan et al., 2018), on previously electroporated embryos at E12.5. As described for *in utero* electroporation, E14.5 pregnant females were anesthetized with isoflurane, analgesic solution was injected, and Caesarean section was performed to expose the uterine horns. 0.5 μL of 10mM FT (CellTraceTM CFSE, Life Technologies, #C34554) was injected into one of the embryos’ lateral ventricles. The utero was then placed in the abdominal cavity, the abdominal wall sutured, and embryos were allowed to develop *in utero* until collection time at E18.5.

### Immunohistochemistry

#### Tissue processing

Embryonic brains were dissected in a phosphate-buffered saline solution (PBS), fixed in 4% paraformaldehyde (PFA) in PBS overnight at 4°C, and later stored in PBS-0.05% sodium azide at 4°C. After embedding in 4% low melting point agarose in PBS, brains were sectioned coronally at 100 μm using a vibratome (Leica VT1200S). Postnatal mice were fixed by transcardial perfusion with 4% PFA before dissection and post-fixation. Coronal sections were performed obtaining 60 μm sections.

#### Immunofluorescence on brain sections

Brain sections were incubated for 1 hour in blocking solution (10% horse serum, 0.2% Triton X-100 in PBS) at room temperature. Sections were incubated overnight with the primary antibody at the appropriated dilution in blocking solution at 4°C. Sections were washed three times with PBS and incubated for 1 hour with the respective secondary antibodies (1:1000) in blocking solution at room temperature. After three washes in PBS, sections were mounted on slides with Fluoromount G (Electron Microscopy Sciences).

#### Antibodies

rat anti-Ctip2 (1:500, Abcam, #AB18465); rabbit anti-Cux1 (1:300, Santa Cruz Biotechnology, sc-13024); rabbit anti-Cux1 (1:1000) (Proteintech, #11733-1-APr); rabbit anti-Doublecortin (DCX) (1:1000, Abcam, #ab18723); goat anti-GFP (1:500, Rockland antibodies & assays, #600-101-215M); rabbit anti-Ki67 (1:500, EMD, #AB9260); rabbit anti-Pax6 (1:500, GeneTex, #GTX113241); mouse anti-Phospho Vimentin (1:500, MBL, D076-3); mouse anti-RFP (1:200, LSBio, LS-C29691); rabbit anti-Satb2 (1:2000, Abcam, #ab92446); rabbit anti-Sox9 (1:1000, Abcam, #AB184547; rabbit anti-Tle4 (1:500, Abcam, #ab140485); rabbit anti-Tbr2 (Eomes) (1:500, Abcam, #AB23345). Fluorescent secondary antibodies were used at 1:1000 (Jackson lmmunoresearch).

#### Imaging

Images were acquired using a Nikon-C2 or a FluoView FV10i (Olympus) confocal laser-scanning microscope. Image analysis and quantification were performed using Photoshop (Adobe), FIJI (ImageJ) and R software.

### Quantification of Tissue Sections

Four different animals were used for most of the conditions with a couple of exceptions where three animals where analysed. Several rostro-caudal sections were selected based on the electroporation location, limiting the use of electroporations to the lateral neocortex including the somatosensory neocortex.

#### Analysis of the developing neocortex at E13.5

Quantifications in the developing neocortex were performed by dividing its surface in two regions: VZ, and outside the VZ, (including the subventricular zone (SVZ), intermediate zone (IZ) and cortical plate (CP)). Density and morphological features were used to differentiate these regions.

#### Analysis of the developing neocortex at E14.5

Quantifications in the developing neocortex were performed by dividing its surface in four regions: VZ, SVZ, IZ and CP. Density and morphological features were used to differentiate these regions.

#### Analysis of the cortical plate at E18.5 and P0

Quantifications at these ages were performed by dividing the cortical plate in ten bins. This bin division was used since the cortical plate is still unmatured and some neurons are still migrating at this stage.

#### Analysis of the cortical plate at P12 and P20

Quantifications of the cortical plate were performed by dividing the neocortex surface in two regions: upper layers (UL) and lower layers (LL). In order to differentiate these two regions, immunochemistry with Cux1 antibody was used to define the UL (II-IV).

#### Analysis of the ventricular zone at E14.5

For every counted cell, position in the ventricle and intensity of fluorescence was measured. Position was calculated by its distance to the ventricle, assigning y=0 to the bottom of the counted area (lateral ventricle) and y=1 to the top (subventricular zone). Intensity was calculated by measuring the pixel intensity in a scale of 0-255 (8-bits). Every cell’s fluorescence intensity was corrected by its particular background value calculated for every counted section:

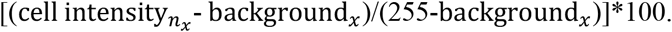

### Tissue Preparation for Single Cell Sorting and Sequencing

Brains electroporated with the pCAG-mRFP;Hes5p-GFP plasmid were dissected 11 h after electroporation and embedded in 4% Agarose in complete HBSS medium (Polleux and Ghosh, 2002) and sectioned using a vibratome (Leica VT1200S). Electroporated areas of the lateral cortex were microdissected under the fluorescent dissecting scope (Zeiss Axio Zoom). Tissue was dissociated into single cells using the Papain Dissociation System (Worthington, #LK0031500). Resuspended cells were processed using a cell sorter (MoFlo XDP) and viable single cells expressing mRFP or mRFP and GFP were individually collected in 96 well plates with lysis buffer. Individual lysed collected cells were processed for scRNAseq using a modified Smart-seq2 protocol (Chevée et al., 2018). Data were aligned to mouse reference genome (mm10) using Hisat2 (Kim et al., 2015) and quantified against a modified reference transcriptome (GENCODE vM15; (Frankish et al., 2018) with cuffquant (Trapnell et al., 2014). Default parameters were set for both software except parallelization value (-p 6) to improve processing time.

### scRNAseq Analysis

Data were analyzed using Seurat R package v2 (Butler et al., 2018). To import data into a Seurat object and eliminate low quality cells/genes (quality-check), CreateSeuratObject function selected cells with at least 300 genes, and genes that were expressed in at least 3 cells. A global-scaling normalization method (LogNormalize by 104 factor) was implemented using NormalizeData. Highly variable genes were detected (FindVariableGenes) across the single cells and data were scaled (ScaleData) to linear transform our data and regress out variables that would not give meaningful information to our analysis such as a high percentage of mitochondrial genes expression and number of molecules detected. RunPCA were used to perform linear dimensional reduction and among those PCAs, 10 (based on PCElbowPlot) was selected to further analysis. Clusters and tSNE 2-dimensional visualization were built using PCAs with FindClusters and RunTSNE functions, respectively. FindAllMarkers were used to identify differentially expressed genes in each group and clusters was renamed based on expression of identified known cell markers.

### Transcriptional Regulatory Networks

For our networks, we used RTN software (Castro et al., 2016) that creates a transcriptional regulatory network and analysis of regulons. As input, scRNAseq data from ventricular zone E13 cells (Yuzwa et al., 2017) were reprocessed and their respective normalized data table were used. To select mouse transcription factors (TFs), a transcription factor database (TcoF-DB v2; (Schmeier et al., 2017) was used to identify TFs in our data. Then, TNI-class object was created using previous described inputs (normalized table and selected TFs) (*tni.constructor*), followed by permutation analysis (*tni.permutation*, nPermutations = 1000) and bootstrap (*tni.bootstrap*). Then, differentially expressed genes from each cluster and their respective average log2 foldchange average expression were used to find master regulons (*tni.mra*). Resulting regulon network were manipulated using RedeR software (Castro et al., 2016)

### Experimental Design and Statistical Analysis

Mean, standard deviation (SD), standard error of the mean (SEM) and statistical analysis were performed using Excel from Microsoft. Statistically significant differences of means were assessed by Student’s unpaired t-test comparing two groups. P-value < 0.05 was considered a significant difference (*). The exacts p value, mean ± SEM and n are reported in the results section.

## RESULTS

### *In Utero* Electroporation Using Different Promoters as a Strategy to Distinguish Progenitors with Different Neurogenic Behavior

It has been reported that a subpopulation of RGCs targeted with constructs using the *Emx2* enhancer/promoter at E12.5 showed delayed neurogenic divisions compared to RGCs targeted with ubiquitous promoters (García-Moreno and Molnár, 2015). We used this strategy to identify progenitors with different behavior and we conducted the experiments at the same embryonic age (E12.5) reported in the earlier study. We selected expression vectors containing promoter fragments driving fluorescent proteins that report the activity of important molecular pathways involved in development of the central nervous system including Wnt signaling (Hirabayashi et al., 2004; Chenn, 2008) and Notch signaling (Yoon and Gaiano, 2005; Louvi and Artavanis-Tsakonas, 2006). We compared the labeling observed using these promoters to that of the general promoter CAG driving BFP (Fig. 1A). To report Wnt signaling we used the Top-GFP plasmid containing a tandem of 7 TCF/LEF promoter elements (Horst et al., 2012). To report Notch activity, we selected a plasmid containing the Hes5 promoter fragment driving dsRed protein expression (Mizutani et al., 2007). We performed *in utero* electroporation using the three plasmids to visualize possible differences in the type of cells activating the different promoters in the same tissue. Twenty-four hours after electroporation using the CAG promoter, we found a considerable number of labeled cells located in the SVZ-Intermediate zone (IZ). A similar result was observed using the Wnt reporter plasmid, showing no significant differences in the percentage of electroporated cells inside the VZ (pCAG-BFP: 47.97 ± 1.23%; TOP-GFP: 43.41 ± 2.45%; *t* (6) = −1.67, *p* = 0.147; unpaired t test, n = 4; Fig. 1B, E). However, with the Hes5 reporter most of the cells were found in the VZ (pCAG-BFP: 50.51 ± 0.66%; Hes5p-dsRed: 81.69 ± 5.47%; *t* (6) = 5.66, *p* = 0.001; unpaired t test, n = 4; Fig. 1B, E). A similar pattern was observed using another Notch reporter plasmid containing the promoter CBF-RE (Mizutani et al., 2007), a tandem of 5 CBF1-binding elements driving the expression of a different fluorescent protein, GFP (pCAG-BFP: 49.67 ± 0.69%; CBFRE-GFP: 83.55 ± 1.91%; *t* (6) = 16.71, *p* = 2.94 × 10^-6^; unpaired t test, n = 4; Fig. 1C, E). Co-electroporation experiments of Hes5p-dsRed and Emx2p-GFP plasmids at E12.5 showed co-localization of the two reporter genes in all of the targeted cells twenty-four hours after electroporation (Fig. 1D-E). Comparison of the distribution of labeled cells using the Emx2 and CAG promoters consistently revealed the existence of statistically significant differences (pCAG-BFP: 50.51 ± 0.66%; Emx2p-GFP: 80.13 ± 4.75%; *t* (6) = 6.17 *p* = 0.00083; unpaired t test, n = 4; Fig. 1D-E). For simplicity, we refer to cells activating the Hes5 promoter as Hes5+ cells, and those activating the CAG promoter as CAG+ cells.

**Figure 1.**
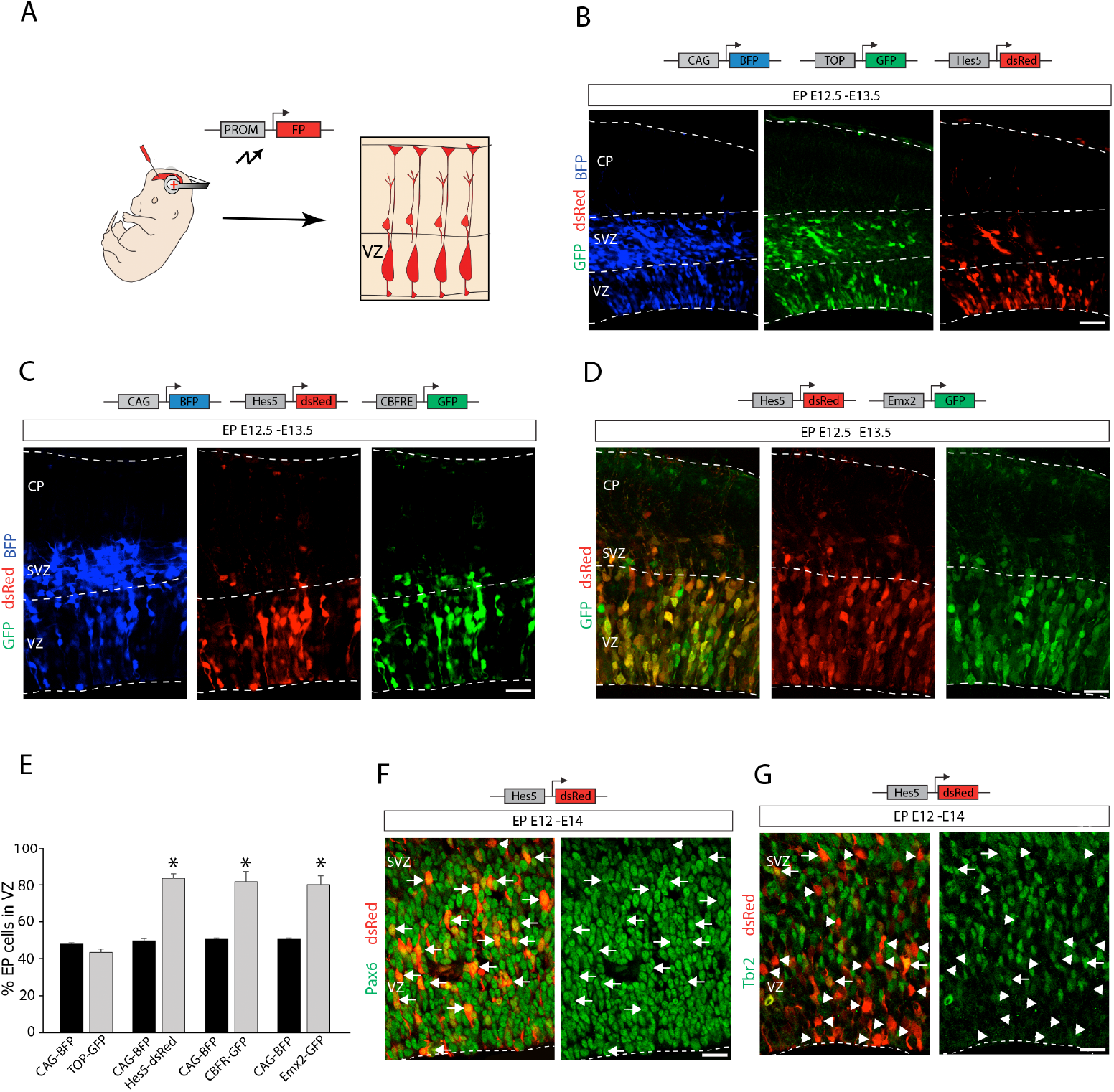
*In utero* electroporation strategy to identify RGC subpopulations. (A) Schematics of the electroporation strategy for labeling different subpopulations of RGCs. PROM, promoter; FP, fluorescence protein. (B) Triple electroporation at E12.5 of pCAG-BFP (blue), Wnt reporter plasmid TOP-GFP (green) and Notch reporter plasmid Hes5p-dsRed (red), analysis at E13.5. Note the similar distribution of labeled cells using the general promoter (pCAG), and the Wnt reporter plasmid showing a notorious number of cells in the SVZ. The Notch reporter plasmid allowed the labeling of cells mostly present in the VZ. (C) Triple electroporation at E12.5 of pCAG-BFP (blue), Notch reporter plasmids Hes5p-dsRed (red), and CBFRE-GFP (green), analysis at 13.5. Note CBFF-RE and Hes5 promoter fragments drive a similar labeling pattern. (D) Co-electroporation of Hes5p-dsRed (red) and Emx2p-GFP (green) at E12.5, analysis at 13.5. Note the virtual co-localization of both reporter proteins in cells activating these promoter fragments. (E) Quantification (mean ± SEM) of electroporated (EP) cells in the VZ at 13.5 with different expression vectors exhibiting differences in their neurogenic behavior. *, *P* <0.002. (F-G) Immunostaining (green) at E14.5 for Pax6 (F) and Tbr2 (G) of E12.5 Hes5-dsRed (red) electroporated embryos. (F) Arrows point examples of dsRed+/Pax6+ cells. (G) Examples of dsRed+/Tbr2+ cells (arrows) and dsRed+/Tbr2-cells (arrowheads). CP, cortical plate; SVZ, subventricular zone; VZ, ventricular zone. Scale bars: 50 μm.

In order to confirm that the Hes5+ cells in the VZ were RGCs, we performed immunohistochemistry. We found that forty-eight hours after electroporation the majority of the Hes5+ cells expressed the RGC marker Pax6 (Fig. 1F, arrows) but not the intermediate progenitor (IP) marker Tbr2 (Fig. 1G, arrow heads).

### Hes5+ RGCs Mostly Generate Upper Layer Cortical Neurons

Hes5+ RGCs at early stages of corticogenesis appear to present delayed neurogenesis compared to other neurogenic progenitors at the same developmental stage. We next analyzed distribution of electroporated cells at later stages after *in utero* electroporation to determine laminar localization of the neurons produced by Hes5+ cells (Hes5p-dsRed) in comparison with CAG+ cells (pCAG-BFP) (Fig. 2A). We analyzed the laminar position of labeled cells seven days after E12.5 electroporation. CAG+ neurons were located within both the upper and lower parts of the developing cortical plate (CP), with greater accumulation in the lower part. On the other hand, Hes5+ neurons mostly accumulated in the upper part of the CP (pCAG-BFP vs Hes5p-dsRed, bin 1: *p* = 2.47 × 10^-5^, bin 2: *p* = 5.52 × 10^-8^, bin 3: *p* = 1.89 × 10^-6^, bin 5: *p* = 0.0002, bin 6: *p* = 0.0106, bin 7: *p* = 5.20 × 10^-6^, bin 8: *p* = 4.60 × 10^-5^, bin 9: *p* = 0.0002, bin 10: *p* = 0.0057, unpaired t test, n = 4; Fig. 2B-C). In order to determine the subtype of neurons that were Hes5+, we conducted immunohistochemistry using known markers for subtypes of neocortical neurons, namely Cux1, Satb2, Tle4 and Ctip2. Cux1 is a specific marker for upper layer neurons (Nieto et al., 2004) and the majority of Hes5+ cells located in the upper part of the CP also expressed Cux1 (85.72 ± 1.35%; Fig. 2D, H). Satb2 is a marker of cortico-cortical neurons, mostly located in upper layers but also present in some lower layer neurons (Alcamo et al., 2008; Britanova et al., 2008). Most of the Hes5+ cells were Satb2+ (85.74 ± 1.22%; Fig. 2E, H). Hes5+ cells did not express Tle4, a marker for cortico-thalamic neurons in layer VI (Koop et al., 1996) or Ctip2, a marker for neurons in layers V and VI (Arlotta et al., 2005) (Hes5+Tle4+/Hes5+: 9.71 ± 1.10%, Hes5+Ctip2+/Hes5+: 5.67 ± 0.98%; Fig. 2F-H). These results suggest that Hes5+ progenitors are involved in the generation of cortico-cortical projection neurons mostly located in upper layers.

**Figure 2.**
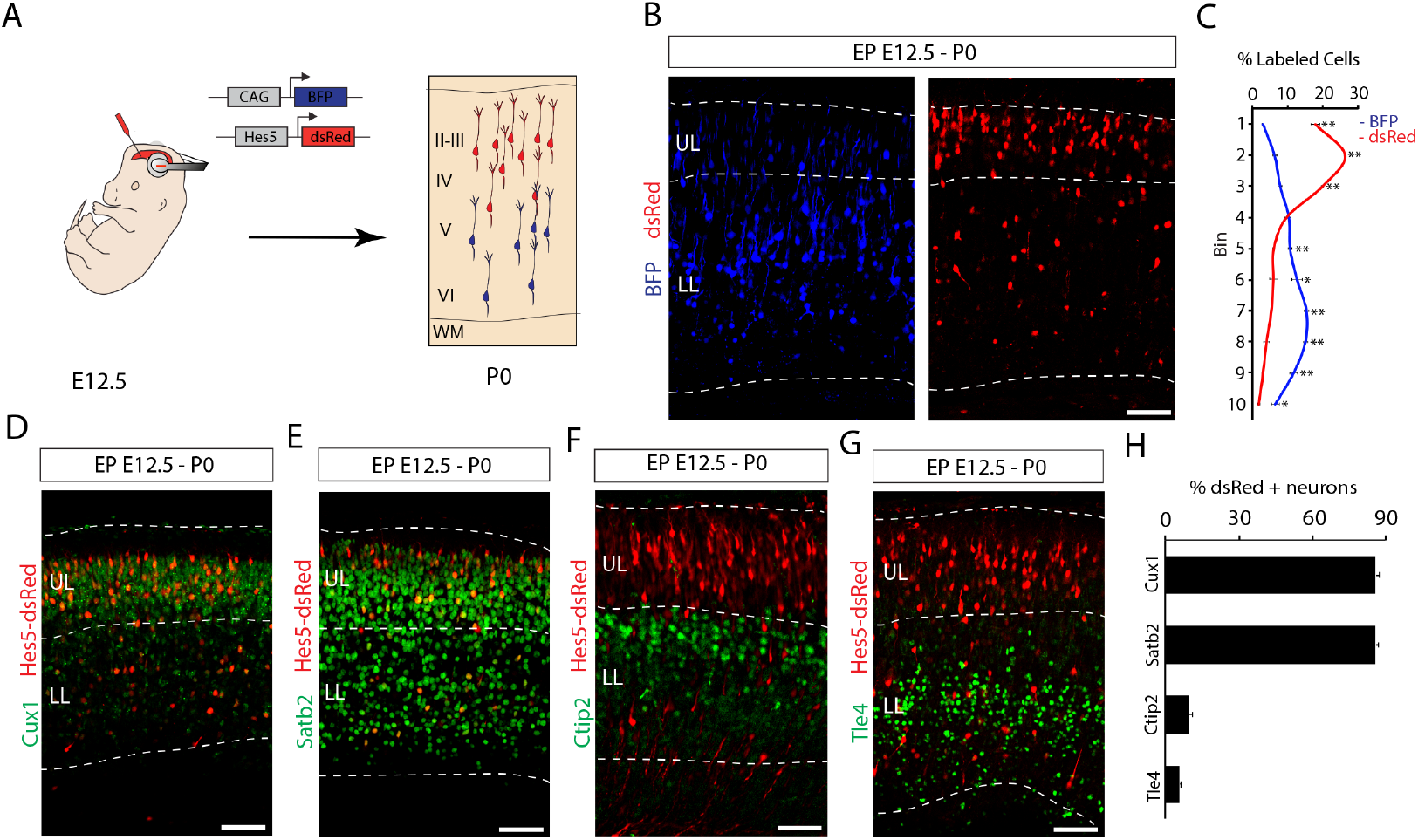
Laminar localization of neurons generated from early RGCs with delayed neurogenic potential. (A) Schematics of the electroporation strategy for laminar localization determination. (B) Co-electroporation of the pCAG-BFP (blue) and Hes5p-dsred (red) constructs at E12.5, analysis at P0. (C) Quantification (mean ± SEM) of the distribution of electroporated cells in the CP at P0 from each subpopulation of progenitors. For quantification, the CP was divided in ten equal-size bins (enumerated 1-10 from basal to apical). *, *P* < 0.02 **, *P* < 0.0005 (D-G) Immunostaining (green) at P0 for Cux1 (D), Satb2 (E), Ctip2 (F) and Tle4 (G), of E12.5 Hes5p-dsRed (red) electroporated embryos. (H) Quantification (mean ± SEM) at P0 of Hes5p-dsRed electroporated cells expressing Cux1, Satb2, Ctip2 or Tle4. II-VI, cortical layers 2-6; WM, white matter; LL, lower layers; UL, upper layers. Scale bars: 100 μm.

Hes5 is not expressed in differentiated neurons (Basak and Taylor, 2007), suggesting that the labeled neurons located in upper layers came from early Hes5+ RGCs. To formally test this possibility, we conducted lineage tracing experiments. We generated a tamoxifen inducible Cre-recombinase construct driven by the Hes5 promoter (Hes5p-CreERT2) to conduct *in utero* electroporation at E12.5 in Ai9 reporter mice (Madisen et al., 2010). As a control, we used a general promoter (pCAG-CreERT2). Since 4-OHT has a short half-life (Guenthner et al., 2013), we injected 4-OHT immediately following *in utero* electroporation to maximize recombination in RGCs taking up the plasmid, before they generated more differentiated cell types. We then analyzed the distribution of the neuronal offspring of electroporated RGCs at P20. Following Hes5p-CreERT2 electroporations and 4-OHT injection, there was a strong bias in the localization of td-Tomato+ neurons towards upper layers when compared to pCAG-CreERT2 electroporations (pCAG-CreERT2 vs Hes5p-CreERT2, bin 1: *p* = 0.0011, bin 2: *p* = 0.0043, bin 3: *p* = 0.002, bin 5: *p* = 0.0037, bin 6: *p* = 0.0417, bin 7: *p* = 0.001, bin 8: *p* = 0.0014, bin 9: *p* = 0.0037, bin 10: *p* = 0.0167, unpaired t test, n = 4; Fig. 3B-C). Most of the td-Tomato+ neurons in brains electroporated with Hes5p-CreERT2 expressed the upper layer marker Cux1 (83.96 ± 7.30%; Fig. 3C, E) and the cortico-cortical marker Satb2 (90.81 ± 0.89%; Fig. 3D-E), even when located in lower layers (Fig. 3D). Very few of them expressed Ctip2 (3.79 ± 1.48%), which is mostly expressed by cortico-fugal neurons and some interneurons (Arlotta et al., 2005) (Fig. 3C-E). In contrast, td-Tomato+ neurons in brains electroporated with pCAG-CreERT2 comprised higher percentages of Ctip2+ neurons (37.97 ± 7.22%; t (4) = 2.13, *p* = 0.0097, unpaired t test, n = 3) and reduced numbers of Cux1+ and Satb2+ recombined neurons (Cux1+:38.70 ±1.98%; t (4) = −5.98, *p* = 0.0039; Satb2+: 56.96 ± 2.34%; t (4) = −13.51, *p* = 0.00017, unpaired t test, n = 3; Fig. 3E).

**Figure 3.**
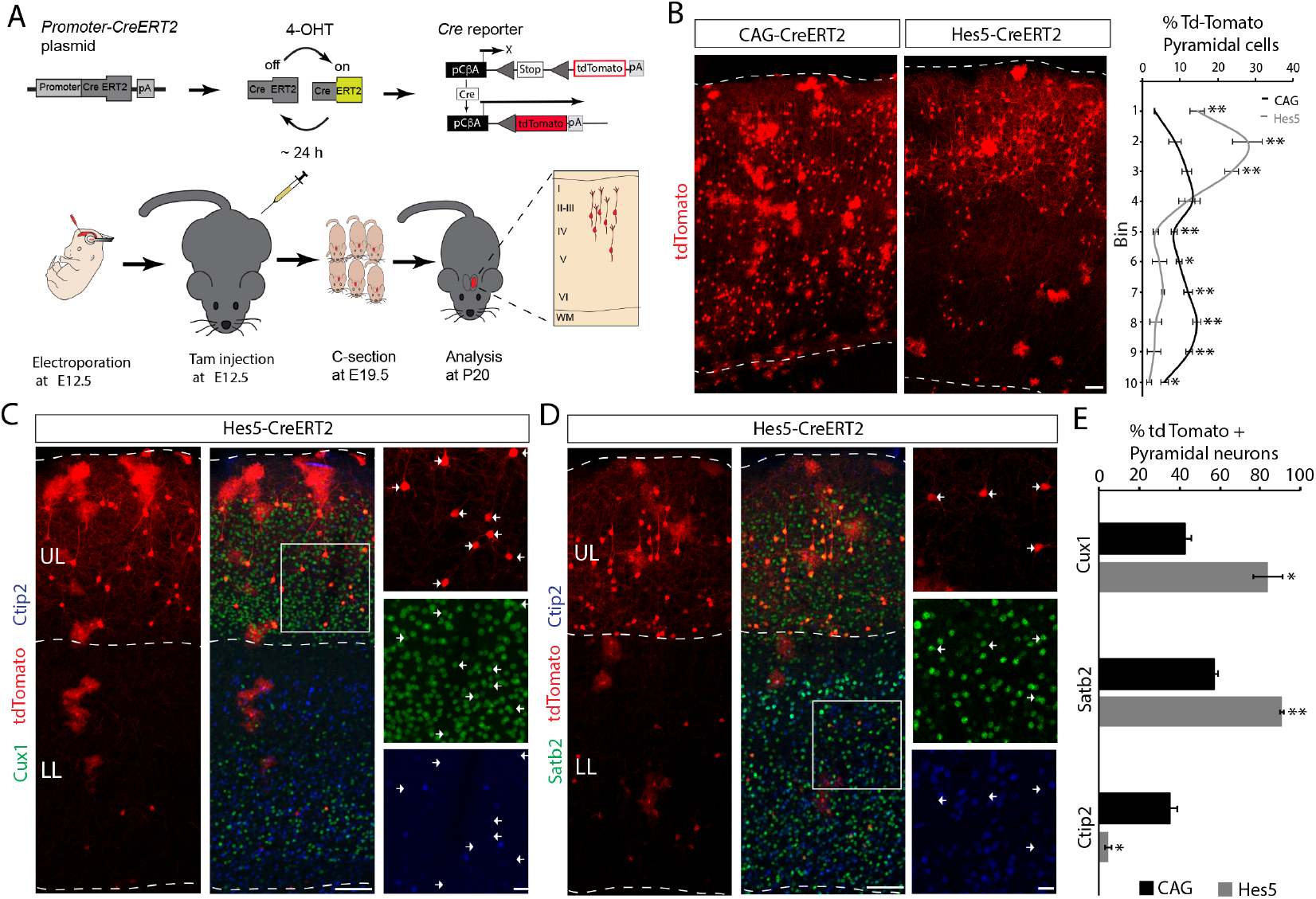
Lineage tracing of early RGCs activating Hes5 promoter. (A) Schematics of the linage tracing strategy. Ai9 reporter embryos were electroporated at E12.5 with pCAG-CreERT2 and Hes5p-CreERT2 constructs. Pregnant dams were injected with 4-OHT right after electroporation. Pups were delivered at E19.5 by Caesarean section. Brains were analyzed at P20. (B) Confocal images and quantification (mean ± SEM) of the distribution of tdTomato+ cells at P20. Note most tdTomato+ cells in the Hes5p-CreERT2 electroporated brains were present in the upper layers, whereas pCAG-CreERT2 show a more homogeneous distribution. For quantification, the CP was divided in ten equal-size bins (enumerated 1-10 from basal to apical). *, *P* < 0.05 **, *P* < 0.005. (C) Double immunofluorescence for Cux1 (green) and Ctip2 (blue) on Hes5p-CreERT2 (red) electroporated brains showing magnification in the upper layers. Arrows point examples of tdTomato+/Cux1+ cells. (D) Double immunofluorescence for Satb2 (green) and Ctip2 (blue) on Hes5p-CreERT2 (red) electroporated brains showing magnification in the lower layers. Arrows point examples of tdTomato+/Satb2+ cells. (E) Quantification (mean ± SEM) on pCAG-CreERT2 and Hes5p-CreERT2 electroporated brains of recombined tdTomato+ cells expressing Cux1, Satb2 or Ctip2. *, *P* < 0.01 **, *P* < 0.0002. Abbreviations as in Figure 2. Scale bars: B 100 μm; C-D 100 μm low magnification and 50 μm boxed areas.

### Single Cell RNA Sequencing of Early Hes5+ RGCs Reveals a Molecular Signature of Cortical Progenitors with Delayed Neurogenic Potential

The previous results suggest the existence of two subtypes of RGCs at early ages: one of them being early neurogenic and other remaining longer as RGCs before generating neurons. To search for molecular differences between the two subtypes of RGCs, we isolated neocortical cells for characterization by single cell RNA sequencing. For this purpose, we carried out *in utero* electroporation with a dual promoter plasmid containing two fluorescent proteins under the control of two different promoters: the CAG promoter driving mRFP and the Hes5 promoter driving GFP. This strategy ensures that both reporter promoters are present within each electroporated cell. We optimized the timing between the surgery for *in utero* electroporation at E12.5 and cell isolation to optimize fluorescence intensity of cells largely confined to the proliferative zone of the neocortex (Fig. 4A). The optimal time point for cell isolation was 16 hours after *in utero* electroporation, with substantial numbers of Pax6+ RGCs (Fig. 4A) expressing either only mRFP (Fig. 4B, arrows) or both fluorescent proteins (Fig. 4B, arrow heads). After brain dissection and isolation of the electroporated areas of the neocortex, we used fluorescence-activated cell sorting (FACS) to separate single-labeled CAG+ cells (only CAG promoter active, mRFP+GFP-cells) and double-labeled CAG+/Hes5+ cells (both promoters active, mRFP+GFP+ cells) (Fig. 4C). Viable sorted cells of both types collected individually were lysed and libraries were generated using a modified Smart-seq 2 protocol for subsequent scRNA sequencing as described (Chevée et al., 2018).

**Figure 4.**
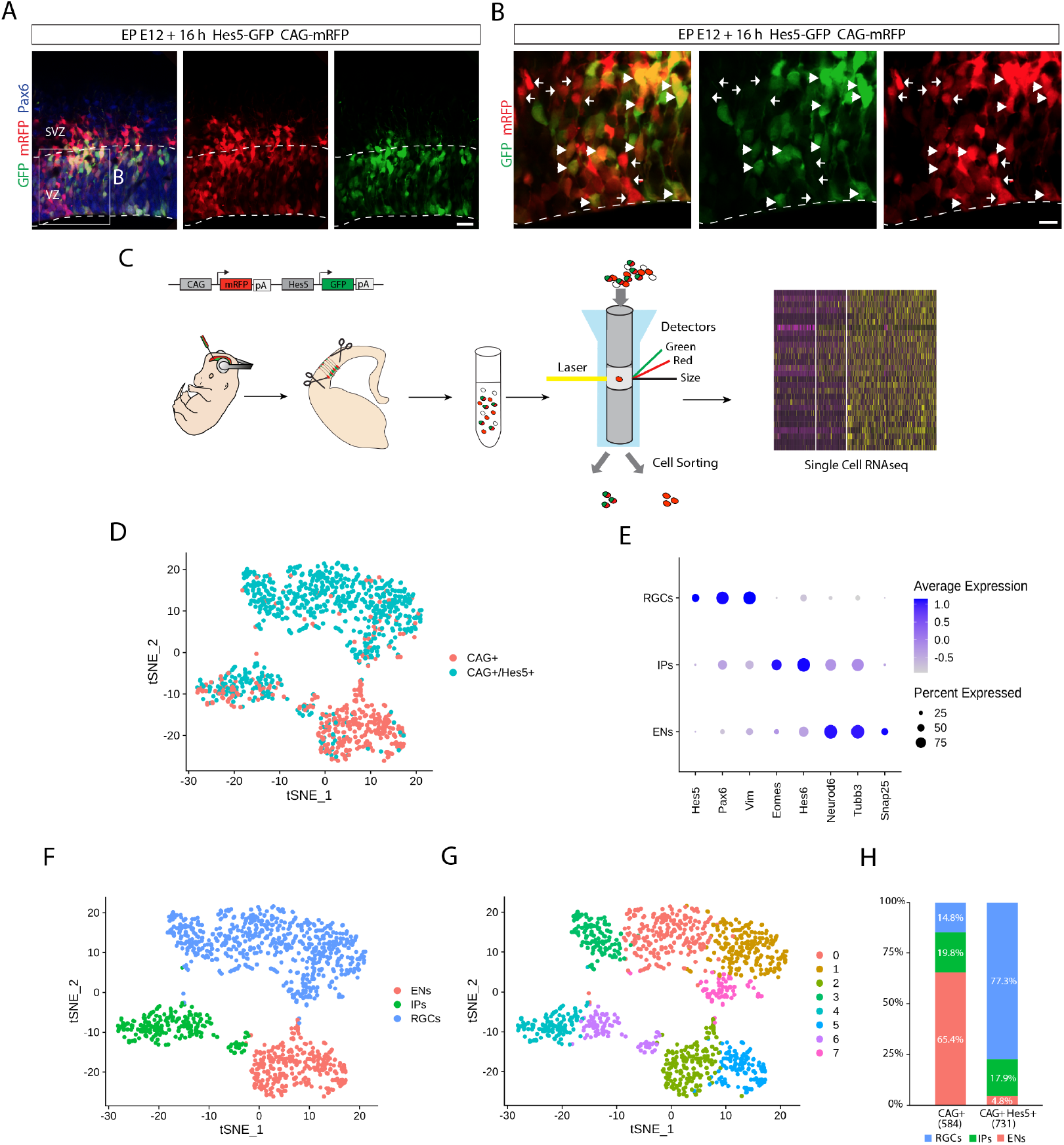
Single-cell sorting, sequencing and scRNAseq identification analysis. (A) Immunostaining for Pax6 (blue) on EP E12.5 electroporated embryos with the dual promotor plasmid pCAG-mRFP; Hes5p-GFP (red and green respectively), analyzed 16 hours after electroporation. (B) Higher-magnification view of boxed area in (A). Arrowheads point mRFP+/GFP+ cells and arrows point mRFP+/GFP-cells. (C) Schematics of scRNAseq strategy. The electroporated area was isolated and cells were individually separated by FACS 16 hours after electroporation. Isolated cells were sequenced and analyzed using bioinformatics tools. (D) t-SNE visualizations of the scRNA-Seq dataset exhibiting mRFP+/GFP+ cell group vs. only mRFP+ cell group. (E) Dot-plot displaying gene average expression (dot gray to dark blue color gradient) and percentage of cells (dot size) expressing cell-type specific markers for RGCs (Pax6, Vim, and Hes5), IPs (Eomes and Hes6), and early neurons (Neurod6, Tubb3, and Snap25) (F-G) t-SNE visualizations of the scRNA-Seq dataset according to expression of the cell-type specific markers (F) and particular clusters identified (G), 0-3 and 7 clusters correspond to RGCs, 4 and 6 clusters are IPs and 2 and 5 clusters are ENs (G). (H) Proportion analysis of Hes5+ and Hes5-cells in the three cell types groups. Hes5+ cells accounted for the majority of cells with an RGC transcriptional signature whereas Hes5-constituted the vast majority of ENs. EN; early neuron; FACS, fluorescence-activated cell sorting; IP, intermediate progenitor; RGC, radial glial cell; SVZ, subventricular zone; VZ, ventricular zone. Scale bars: A. 50 μm and B. 25 μm.

Data were analyzed using the Seurat R package (Butler et al., 2018). A total of 1115 cells passed our quality control test. From those cells, we obtained 731 CAG+/Hes5+ (mRFP+ and GFP+) and of 384 CAG+ cells (mRFP+ but GFP-). Transcriptional analysis among single cells identified cell clusters characterized by similar gene expression patterns (Fig. 4D). By using celltype specific markers, these clusters could be assigned to known cell types of the neocortex (Fig. 4E-F). Gene expression of cell-type specific markers for RGCs (Pax6, Vim, Hes5) (Schnitzer et al., 1981; Götz et al., 1998; Hatakeyama et al., 2004), IPs (Eomes, Hes6) (Englund et al., 2005; Kawaguchi et al., 2008), and neurons (Neurod6, Tubb3, Snap25) (Caccamo et al., 1989; Oyler et al., 1989; Goebbels et al., 2006) allowed us to assign the three cell types to different clusters (Fig. 4F-G). According to marker expression, clusters 0, 1, 3 and 7 were RGCs, clusters 4 and 6 IPCs, and clusters 2 and 5 early differentiating neurons (ENs) (Fig. 4G).

We next analyzed how the Hes5+ (mRFP+GFP+) and Hes5-(mRFP+GFP-) cells were distributed between the different cell-type clusters (Fig. 4H). Whereas the proportion of cells that were IPs was similar for both groups (17.9 % of GFP+ cells and 19.8% of GFP-cells), most of the Hes5-cells were categorized as ENs (65.3% of GFP-cells), but only small percentage of the Hes5+ cells were neurons (4.8 % of GFP+ cells), suggesting that Hes5-RGCs are fast neurogenic at this embryonic stage. Conversely, Hes5+ cells largely had the transcriptional signature of RGCs (77.3% of GFP+ cells), while only a minor fraction of Hes5-cells displayed RGC characteristics (14.8% of GFP-cells).

From the identified RGCs we compared the transcriptome of both groups and identified 11 genes whose expression was significantly upregulated in the Hes5+ subpopulation of RGCs (adjusted p-value < 0.05 and average log2FC > 0.3, Fig. 5A), suggesting possible roles in the regulation of the different cellular behaviors of the two populations of RGCs. In order to investigate important regulators in this event, we generated a transcriptional regulatory network and searched for master regulons (Margolin et al., 2006; Fletcher et al., 2013; Castro et al., 2016). A regulatory network was built using previously published single cell RNA sequencing data at the same age as our analysis (E13) (Yuzwa et al., 2017) (total of 1944 cells and 16378 genes). We found a regulatory network with 49 regulons, which shares commonly regulated genes based on proximity of their neighbor regulons (Fig. 5B). Sox9 and Hes5 regulons were next to each other in our network, which indicates a similar regulatory mechanism between them. Using our network and differentially expressed genes from each cell cluster (Hes5+ or Hes5-RGCs, IPs and ENs, total of 6 comparisons) also helped us to identify Sox9 as a putative master regulator specific to the Hes5+ RGCs (Fig. 5B).

**Figure 5.**
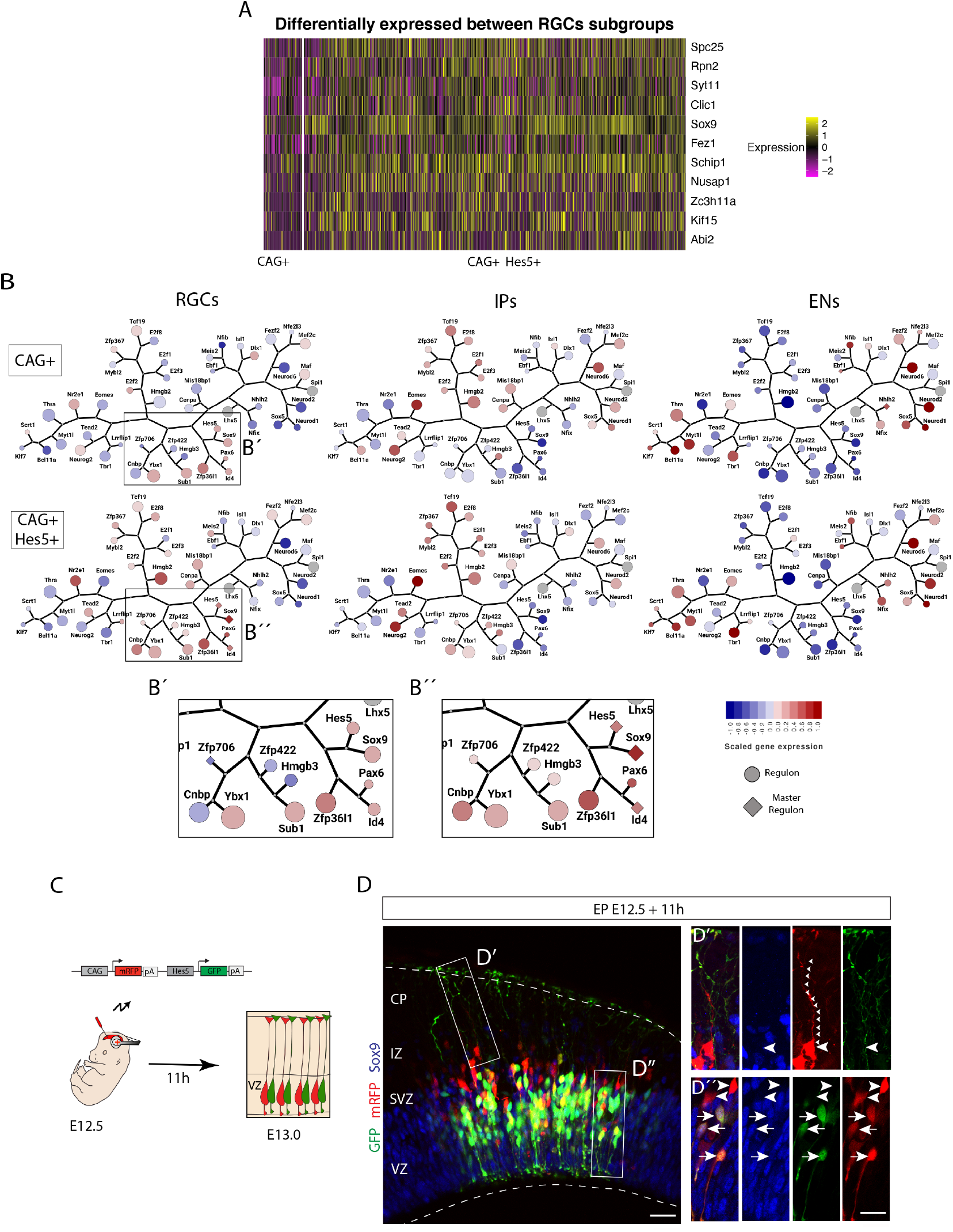
Transcriptional analysis reveals differentially expressed genes and master regulons. (A) Heatmap of differentially expressed genes (rows) in RGCs subgroups comparison (Hes5+ vs. Hes5-) (columns). Gene expression levels are colored ranging from two-fold downregulation (pink) to two-fold upregulation (yellow), according to the color key. (B) Transcriptional regulatory network (regulons) of scRNA-Seq data exhibits Sox9 as a master regulator gene. Circles are regulons and diamond master regulons. Circle size is related to number of genes orchestrated by its transcription factor. Gene expression levels are colored according to average expression of each group. Tree network represent correlation among regulons. (B’-B”) Higher-magnification visualizations of transcriptional network on B boxed areas in Hes5-RGCs (B’) and Hes5+ RGCs (B’’). (C) Schematic illustration of the electroporation strategy to verify Sox9 expression differences indicated by scRNA-Seq analysis. E12.5 embryos were electroporated with the dual promotor plasmid pCAG-mRFP;Hes5p-GFP (red and green, respectively), analyzed 11h after electroporation. (D) Immunostaining for Sox9 after pCAG-mRFP;Hes5p-GFP electroporation. (D’-D’’) Higher-magnification of D boxed areas. Examples of Hes5-RGCs (arrowheads) and Hes5+ RGCs (arrows). Note that Hes5+ RGCs show higher levels of Sox9 immunostaining. Small arrowheads point to the basal process of an RGC. CP, cortical plate; EN; early neuron; IP, intermediate progenitor; IZ, intermediate zone; RGC, radial glial cell; SVZ, subventricular zone; VZ, ventricular zone. Scale bars: D. 50 μm and D’-D” 20 μm.

To verify differences in Sox9 protein expression between the two RGCs subpopulations we performed *in utero* electroporation using the dual promoter plasmid and conducted Sox9 immunohistochemistry 11 hours after the surgery (Fig. 5C). Analysis of Sox9 staining in Hes5+ (Fig. 5D, arrows) and Hes5-(Fig. 5D, arrow heads) RGCs confirmed differences in its expression levels between the two subpopulations.

### Sox9 Overexpression in RGCs Affects Early Neuronal Differentiation

Sox9 is required for stem cell maintenance in different tissues (Scott et al., 2010; Roche et al., 2015) and suppresses cell differentiation (Kadaja et al., 2014). In the CNS, overexpression of Sox9 in the adult subventricular zone (aSVZ) abolishes the production of neurons, whereas Sox9 knockdown increased neurogenesis and reduced gliogenesis (Cheng et al., 2009). We predicted that increased levels of Sox9 observed in Hes5+ progenitors may affect the switch between non-neurogenic and neurogenic cell divisions. To test this hypothesis, we overexpressed Sox9 in RGCs at E12.5 using a general promoter driving Sox9 and GFP and analyzed possible changes in differentiation at E14.5 (Fig. 6A). Unlike control cells expressing only GFP (pCIG), cells overexpressing Sox9 and GFP did not invade the CP (Fig. 6B-C) and accumulated more pronouncedly in the VZ (pCIG: 21.18 ± 1.1%; pCIG-FLS9: 37.67 ± 2.16%; *t* (6) = 6.80, *p* = 0.0005; unpaired t test, n = 4; Fig. 6B,D). Overexpression of Sox9 did not noticeably affect the RGCs scaffold as revealed by the visualization of their end feet (Fig. 6C). Detailed analysis of the cells accumulated in the VZ using markers for RGCs (Pax6), IPs (Tbr2) and postmitotic neurons (Dcx) revealed that the Sox9 overexpressing cells were RGCs (Fig. 6E-G). These results suggest that high levels of Sox9 in RGCs cause a delay in neuron production during early stages of neurogenesis.

**Figure 6.**
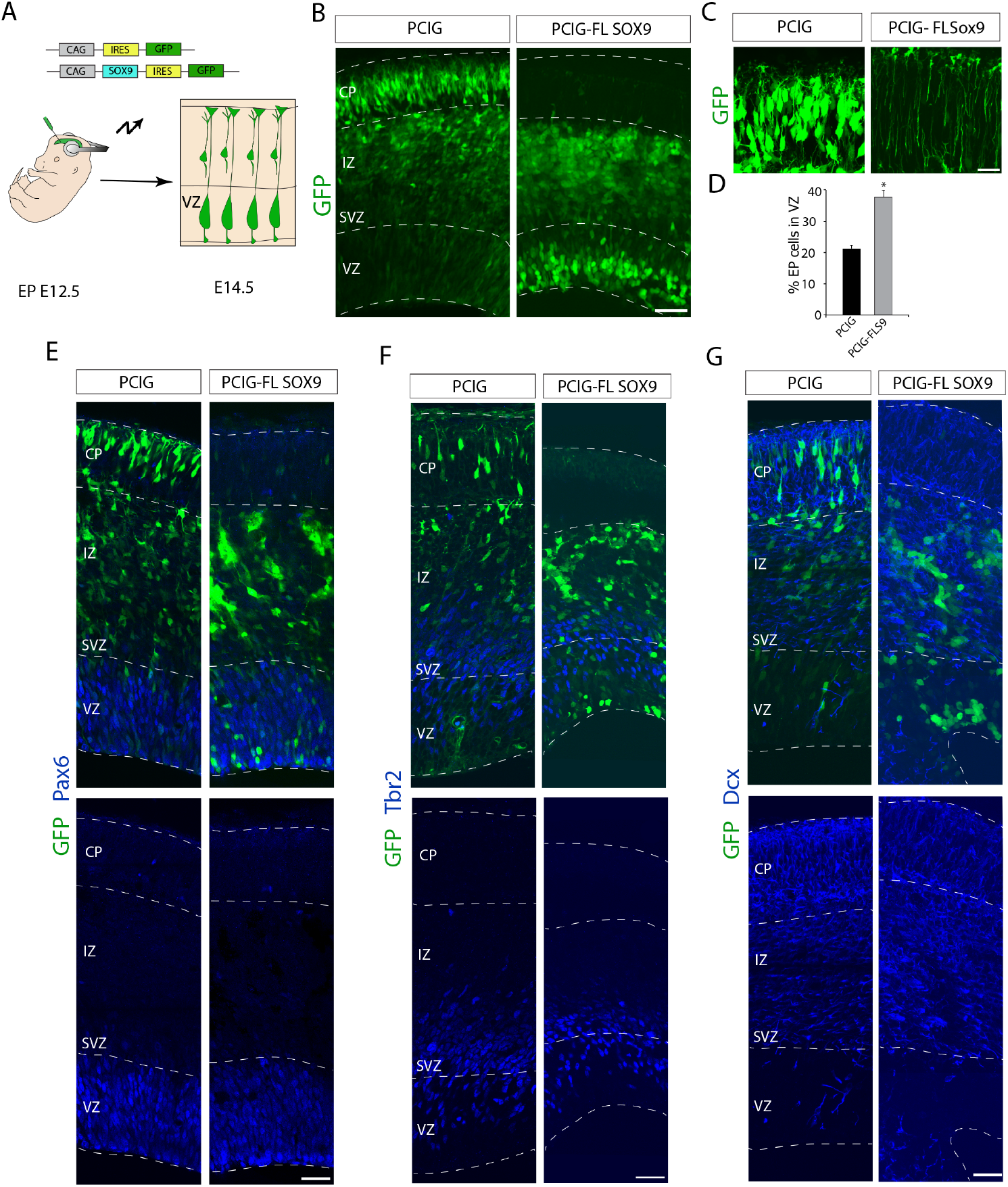
Sox9 overexpression affects neurogenic behavior of RGCs. (A) Schematics of the electroporation strategy. E12.5 embryos were electroporated with the pCIG-FL Sox9 plasmid, to overexpress Sox9, and its empty version, the pCIG plasmid (GFP alone), was used as control. Brains were analyzed 48 hours after electroporation. (B) Confocal images of E14.5 embryo’s neocortex after pCIG or pCIG-FL Sox9 electroporation (green). Note that Sox9 overexpression caused a lack of neurons in the CP and cellular accumulation in the VZ compared to the control condition. (C) Higher-magnification image of the CP showing RGCs’ end feet. Note that Sox9 overexpression did not affect the RGC scaffold. (D) Quantification (mean ± SEM) of targeted cells in the ventricular zone in both conditions exhibits an increase in the Sox9 overexpression situation. *, *P* < 0.0005. (E-G) Immunochemistry at E14.5 brains for Pax6 (E), Tbr2 (F) and Dcx (G) (blue) on E12.5 pCIG or pCIG-FL Sox9 (green) electroporated embryos. n = 4 quantified animals per experiment. CP, cortical plate; IZ, intermediate zone; SVZ, subventricular zone; VZ, ventricular zone. Scale bars: B. 100 μm; C. 20 μm; E-G 50 μm.

### Sox9 Overexpression Acts by Keeping RGCs in a More Quiescent State

Sox9 is associated with quiescence and stem cell maintenance in different tissues including the intestine (Roche et al., 2015) and the hair follicles (Kadaja et al., 2014). In hair follicles, loss of Sox9 results in decreased maintenance of quiescent stem cells with loss of hair follicle regeneration (Kadaja et al., 2014). To determine whether elevated levels of Sox9 in cortical RGCs at early embryonic ages promote progenitor cell quiescence, we carried out further analysis. In *in utero* electroporation experiments, fluorescence intensity of the episomal plasmids is inversely correlated with the number of cell divisions, since the plasmid dosage is diluted in each cell division (Fig. 7A). Only cells exiting the cell cycle quickly after electroporation (postmitotic neurons) or RGCs undergoing limited numbers of cell divisions will maintain strong levels of fluorescent protein expressed from episomal plasmids. Visual inspection suggested that cells overexpressing Sox9 and GFP displayed increased GFP fluorescence compared to cells expressing GFP only (Fig. 7B). From immunohistochemistry analysis at the same age we know that those Sox9 overexpressing cells expressed the RGC marker Pax6, revealing their progenitor nature (Fig. 6E). Quantification of the GFP intensity values in the VZ confirmed statistically significant differences in GFP levels in Sox9 overexpressing cells compared to controls (inner VZ: pCIG: 24.13 ± 2.56; pCIG-FLS9: 91.99 ± 8.59; *t* (6) = 7.57, *p* = 0.0003; outer VZ: pCIG: 27.01 ± 2.83; pCIG-FLS9: 52.34 ± 5.35; *t* (6) = 4.19, *p* = 0.0058, unpaired t test, n = 4; Fig. 7C). The number of Sox9 overexpressing cells in the inner and outer part of the VZ was increased compared to controls, presenting the most obvious difference in the inner part of the VZ (inner VZ: pCIG: 46.59 ± 2.55%; pCIG-FLS9: 65.37 ± 1.63%; *t* (6) = 6.20, *p* = 0.00081; outer VZ: pCIG: 53.41 ± 2.55%; pCIG-FLS9: 34.63 ± 1.63%; *t* (6) = −6.20, *p* = 0.00081, unpaired t-test, n = 4; Fig. 7D). We next co-analyzed GFP fluorescence intensity and distance to the ventricle for individual cells (Fig. 7E). Notably, this co-analysis revealed a remarkable difference between Sox9 overexpressing cells and controls, with brightly labeled Sox9 overexpressing cells enriched near the ventricle surface.

**Figure 7.**
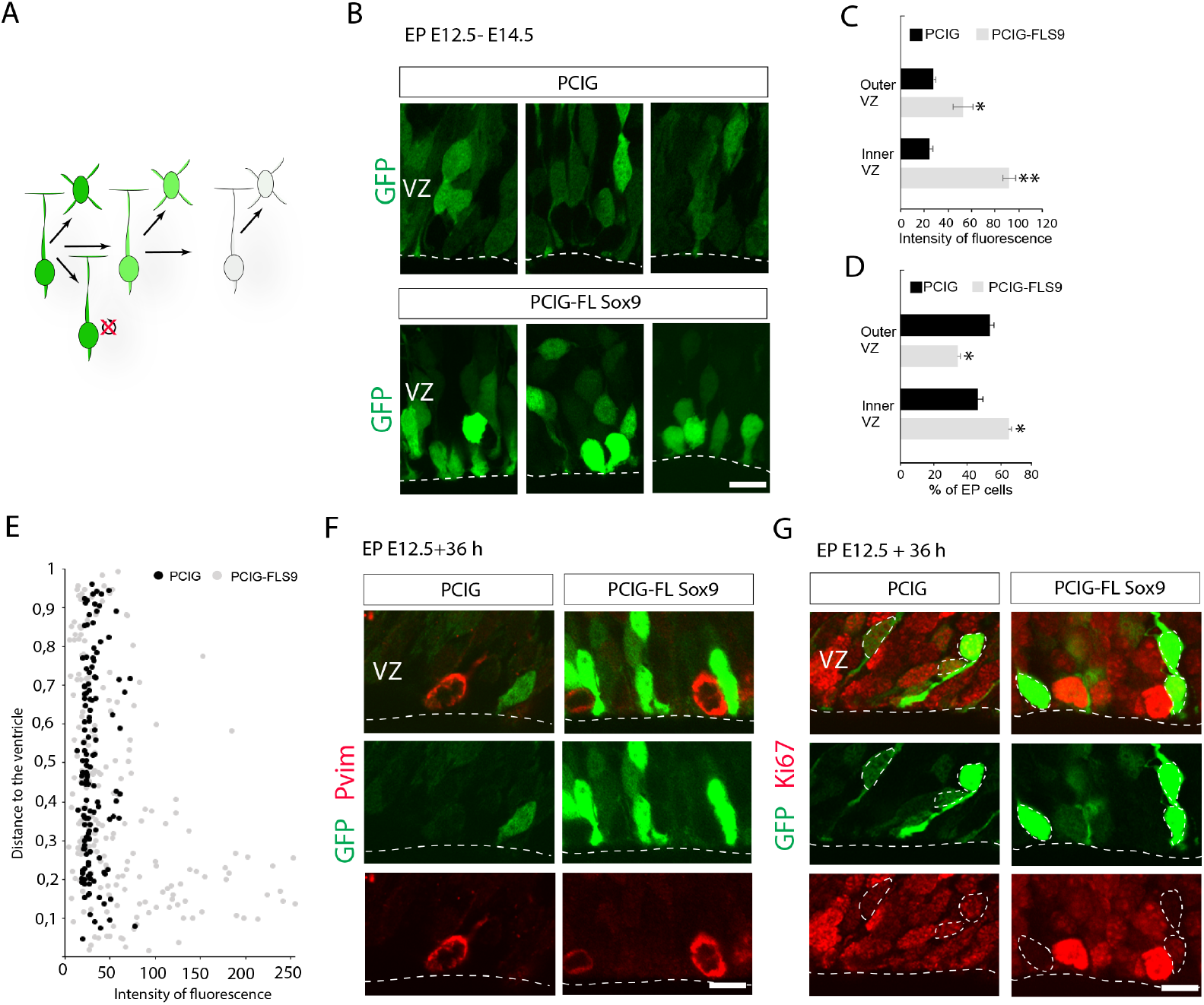
RGCs overexpressing Sox9 display more quiescent behavior. (A) Schematics of the inverse relationship between fluorescence intensity after episomal vector electroporation and cell divisions. (B) High-magnification view of the VZ of pCIG or pCIG-FL Sox9 (green) electroporated brains. Note the increased fluorescence intensity of RGCs after Sox9 overexpression. (C-D) Quantification (mean ± SEM) of the intensity of fluorescence (C) and the distribution (D) of electroporated cells within the VZ in both conditions. For quantification, the VZ was divided in two equal-sized regions, the inner and outer VZ. (C) *, *P* < 0.01 **, *P* < 0.0005; (D) *, *P* < 0.001. (E) Example of co-analysis of fluorescence intensity and distance to the ventricle in the VZ. Every dot represents both parameters obtained from a single cell. Note that Sox9 overexpression condition exhibits a similar dot distribution to the control but additionally displays an accumulation of bright electroporated cells close to the ventricle’s surface. (F-G) Immunochemistry for M-phase marker pVim (F) and proliferation marker Ki67 (G) (red), 36 hours after E12.5 pCIG or pCIG-FL Sox9 electroporation (green). Note that bright electroporated cells in the ventricle’s surface in the Sox9 overexpression condition do not colocalize with pVim (F) and show lower levels of Ki67 than cells expressing GFP alone (pCIG). In (C) and (E) fluorescence intensity was measured in a scale from 0-255. In (E) distance to the ventricle has been relativized to 1. VZ, ventricular zone. Scale bars: 10 μm.

To further analyze the cell cycle in progenitor cells, we performed immunohistochemistry for the M-phase marker phosphorylated vimentin (pVim) (Kamei et al., 1998; Noctor et al., 2002) and the cell cycle marker Ki67 that stains proliferating cells more generally (Gerdes et al., 1983, 1984). The experiments were carried out 36 h after *in utero* electroporation at E12.5 to overexpress Sox9. As shown above (Fig. 7D), the number of GFP labeled RGCs in the inner VZ was increased following Sox9 overexpression compared to controls. These cells did not express significant amounts of pVim (Fig. 7F) and low levels of Ki67 (Fig. 7G) compared to control cells, suggesting that RGCs overexpressing Sox9 are in a more quiescent state compared to other RGCs.

### Sox9 Overexpressing RGCs Engage in Neurogenic Divisions at Later Stages of Cortical Development

Our data suggest that Sox9 levels affect RGC proliferation and their neurogenic potential, such that RGCs expressing high levels of Sox9 are maintained as RGCs during early stages of neocortical neurogenesis. We hypothesized that these cells would generate neurons at later stages of neurogenesis. To test this hypothesis, we designed an experiment to target the RGCs *in vivo* at two different embryonic stages. We first carried out *in utero* electroporation at E12.5 to coexpress Sox9 and GFP or only GFP in RGCs. Next, at E14.5 we performed an intraventricular injection of a carboxyfluorescein ester also known as FlashTag (Telley et al., 2016; Govindan et al., 2018) (Fig. 8A). FlashTag injection will labels RGCs lining the lateral ventricle at E14.5, including those RGCs targeted at E12.5 by in utero electroporation that are still in contact with the ventricle. Thus, cells labeled with both GFP and FlashTag are progenitors that were in contact with the ventricle at both E12.5 and E14.5, whereas GFP+ cells that are not labeled with FlashTag had left the VZ by E14.5. We then analyzed cell distribution of the offspring at E18.5. In controls, most GFP+ cells were FlashTag-negative and accumulated in the lower part of the CP, while FlashTag+/GFP-cells (blue) were largely located in the upper CP (Fig. 8B-C). Following Sox9 overexpression, a substantial number of GFP+ cells occupied the upper CP, the same area also occupied by FlashTag+ cells (Fig. 8C-D). Analysis of single optical sections confirmed coexpression of GFP and FlashTag in these cells (pCIG FT+ in UL: 4.09 ± 0.47 %, pCIG-FLS9 FT+ in UL: 36.36 ± 4.17 %; t (4) = −7.69, *p* = 0.0015, unpaired t test, n = 3; Fig. 8B, D-E) confirming that those neurons were generated from RGCs targeted at E12.5 and still present in the VZ at E14.5. We thus conclude that RGCs expressing high levels of Sox9 are maintained longer at the VZ before undergoing neurogenic divisions.

**Figure 8.**
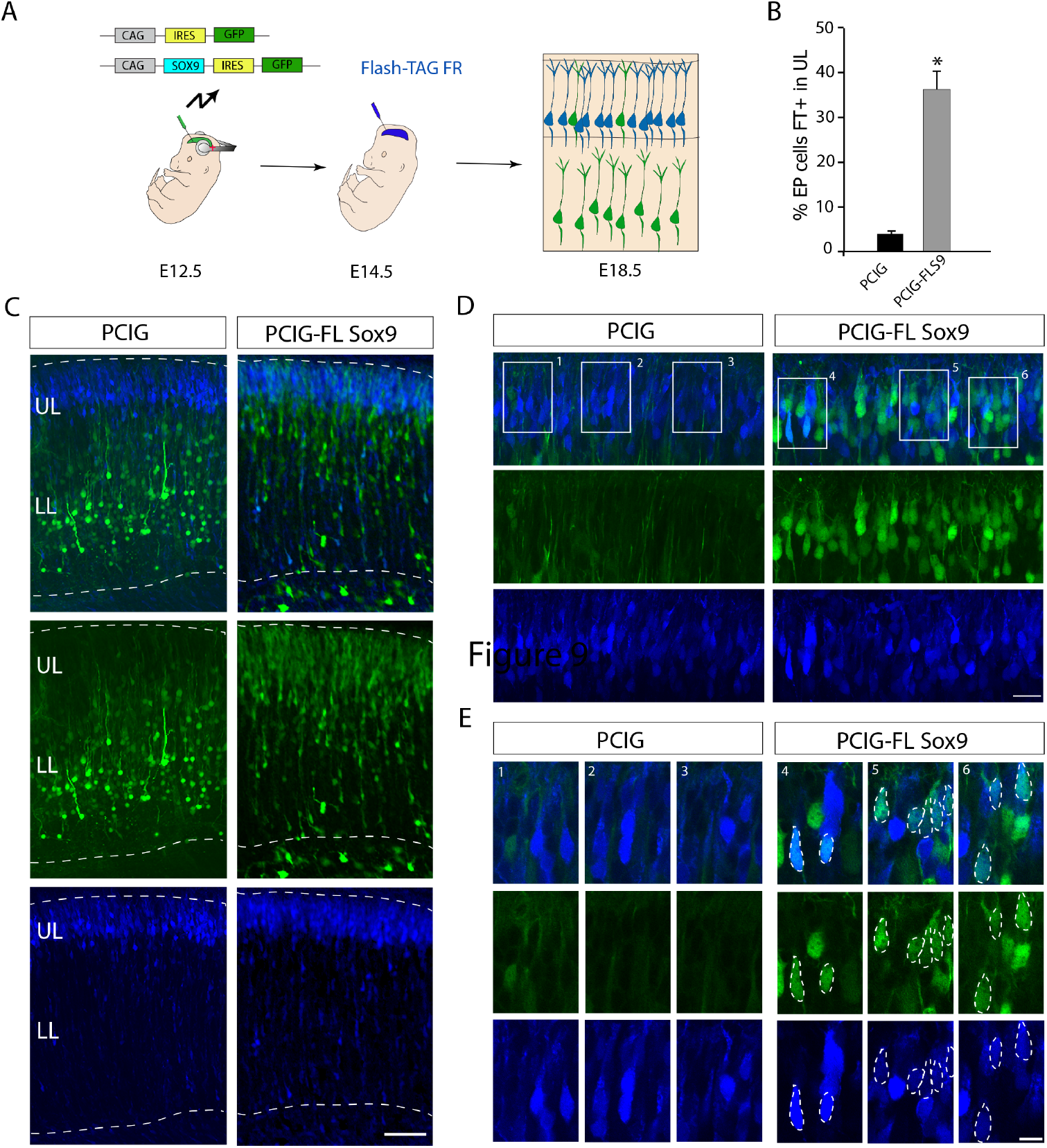
Sox9 overexpressing RGCs engage in neurogenic division at later ages. (A) Schematics of the experimental strategy combining electroporation and FlashTag technologies. E12.5 embryos were electroporated with the pCIG or pCIG-FL Sox9 plasmids (green). 48 hours after electroporation embryos were injected with FlashTag (blue). Brains were analyzed at E18.5. (B) Quantification (mean ± SEM) of GFP+/FT+ double labeled cells in the CP at E18.5 in both conditions. *, *P* < 0.002. (C) Confocal image of E18.5 brains after electroporation and FT injection at described timepoints. (D) Higher-magnification view of (C) showing the FT labeled cells area. (E) Higher-magnification view of (D) boxed areas. Dotted outline points examples of GFP+/FT+ double labeled cells. LL, lower layers; UL, upper layers. Scale bars: C. 100 μm; D. 20 μm; E. 10 μm.

### Sox9 Overexpressing RGCs Produce Neurons Destined for Upper Cortical Layers

To further investigate the fate of neurons derived from RGCs with high Sox9 expression levels, we quantified their laminar positions. We carried out *in utero* electroporation at E12.5 to overexpress Sox9 with GFP or GFP alone and determined the laminar position of the neuronal offspring of electroporated RGCs at E18.5 and P12. Following Sox9 overexpression, more neurons occupied the upper part of the CP compared to controls (pCIG vs pCIG-FLS9, bin 1: *p* = 0.0079, bin 2: *p* = 0.0111, bin 3: *p* = 0.0008, bin 7: *p* = 0.0018, bin 8: *p* = 0.0008, bin 9: *p* = 0.0002, bin 10: *p* = 0.0001, unpaired t test, n = 4; Fig. 9B-C). At P12, when cell migration is complete, these neurons occupied upper neocortical cell layers (pCIG: 26.14 ± 3.14%, pCIG-FLS9: 44.1 ± 4.48%; *t* (6) = 3.28, *p* = 0.0167, unpaired t test, n = 4; Fig. 9D-E). These results are consistent with the model that the cortical VZ contains a subset of RGCs expressing high levels of Sox9, which are maintained as progenitors during early stages of neurogenesis and generate later on upper layer neurons.

**Figure 9.**
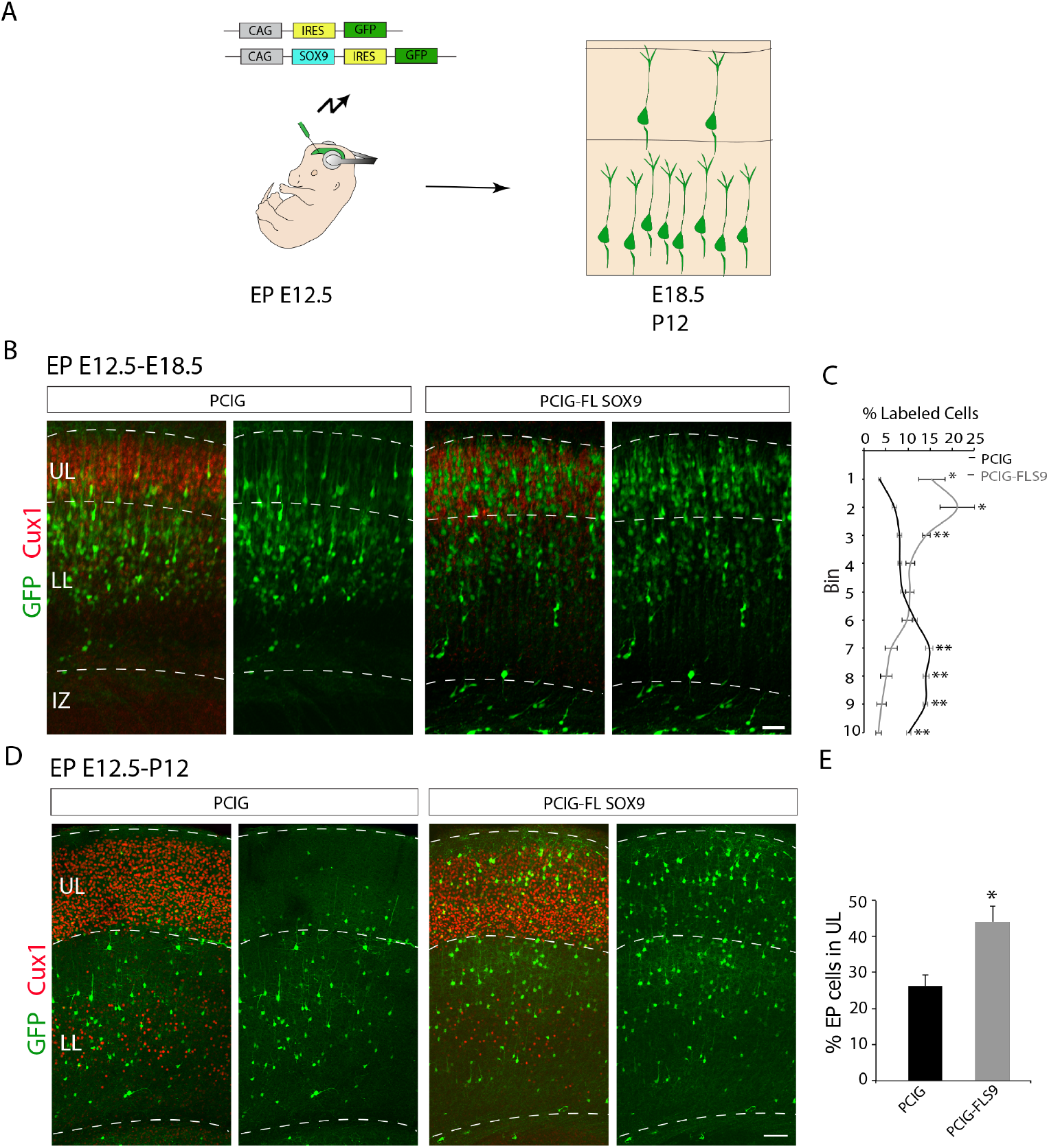
Increased upper layer production upon Sox9 overexpression at early ages. (A) Illustration of electroporation strategy. E12.5 embryos were electroporated with pCIG or pCIG-FL Sox9 constructs (green). Brains were analyzed at E18.5 or P12. (B) Immunostaining for Cux1 (red) on E18.5 brains after E12.5 electroporation of both conditions (green). (C) Quantification (mean ± SEM) of the distribution of electroporated cells within the CP at E18.5. For quantification, the CP was divided in ten equal-size bins (enumerated 1-10 from basal to apical). *, *P* < 0.02, **, *P* < 0.002. (D) Immunostaining for Cux1 (red) on P12 postnatal brains after E12.5 electroporation of both conditions (green). (E) Quantification (mean ± SEM) of the distribution of electroporated cells within the CP at P12. For quantification, the CP was divided in two areas, the lower and upper layers of the neocortex. *, *P* < 0,02. IZ, intermediate zone; LL, lower layers; UL, upper layers. Scale bars: 100 μm.

## DISCUSSION

We provide here evidence of the existence of molecular differences in the pool of neocortical RGCs. Specifically, we identified the molecular signature of a subpopulation of RGCs presenting a more delayed neurogenic behavior at early embryonic ages than other progenitor cells. From the differentially expressed genes between the two populations we identified Sox9 as a specific master regulator of these progenitors. Our functional experiments modifying Sox9 expression revealed that Sox9 levels play a critical role in regulating the decision of neocortical progenitors to be maintained as RGCs or to undergo neurogenic divisions. Our data suggest that high levels of Sox9 maintain RGCs in a relatively quiescent stage, thus promoting their self-renewal and preventing neurogenic divisions during early stages of neurogenesis, to then generate neurons during later stages of neocortical development.

Roles for Sox9 in the control of neuronal stem cells maintenance and differentiation have already been described in several parts of the brain, including the cerebellum (Vong et al., 2015) and the postnatal subventricular zone (Cheng et al., 2009). Our data indicate that Sox9 is important in the regulation of neural stem cell behavior in the early embryonic neocortex. Interestingly, Sox9 is expressed widely in all RGCs, and lineage-tracing experiments using Sox9-CreERT2 mice showed that neurons of all neocortical cell layers were labeled (Kaplan et al., 2017). These data suggest that the levels of Sox9 in RGCs are likely more important than presence or absence. Indeed, a similar dose-dependent regulation of RGC neurogenic behavior has been described for Sox2, another transcription factor of the same family as Sox9 (Hagey and Muhr, 2014). Higher levels of Sox2 have been found in slow-cycling RGCs, and Sox2 overexpression by *in utero* electroporation reduces their proliferation rate.

In the adult subventricular zone, increased Sox9 levels are a marker of quiescent NSCs (Llorens-Bobadilla et al., 2015). However, these progenitors need to be activated to generate neurons at appropriate times, similar to the activation of VZ progenitors to switch from a selfrenewal state during early phases of corticogenesis to neurogenic divisions during late stages of cortical development. How the neurogenic brake is overcome, and which molecular pathways are involved is unclear. One interesting molecular pathway is the Yap/Hippo pathway. Hippo signaling has been related with tissue homeostasis, organ size and stem cell self-renewal (Pan, 2010; Schlegelmilch et al., 2011; Zanconato et al., 2016). A recent report has described that Yap1/Taz promotes NSC maintenance in the cortex and the production of cortico-cortical neurons at the expense of corticofugal neurons (Mukhtar et al., 2020). Interestingly, Sox9 has been described as a downstream target of Yap1 signaling and Yap1-targeting microRNAs, induced by Sox9, post-transcriptionally repress Yap1 expression, contributing to a negative feedback loop (Wang et al., 2019). Yap1 expression is high at early corticogenesis stages and its expression decreases at mid corticogenesis (E14.-E16.5) to increase again at early postnatal ages (Mukhtar et al., 2020). It will be interesting to test in the future whether the decrease in Yap1 expression during mid neurogenesis is critical for determining Sox9 levels and the neurogenic potential of RGCs.

The extent in which different subpopulations of RGCs contribute directly to the generation of cellular diversity in the neocortex is a controversial topic (Franco et al., 2012; Fuentealba et al., 2015; García-Moreno and Molnár, 2015; Gil-Sanz et al., 2015). The results shown here reveal the existence of molecular heterogeneity in RGCs associated with the production of particular cell types. A previously published scRNA sequencing study did not detect such molecular diversity (Telley et al., 2019). However, this study used a different method to label and isolate RGCs; instead of *in utero* electroporation, they used FlashTag technology. Notably, FlashTag labels RGCs with large endfeet at the VZ, which are large dividing cells in M-phase (Govindan et al., 2018). At early embryonic ages (E12.5), the progeny of these labeled cells rapidly exits the VZ to produce IPs or ENs (Telley et al., 2016, 2019). Thus, 10 hours after FlashTag labeling as conducted in this study, most of the labeled cells were found in the SVZ-IZ. In contrast, our *in utero* electroporation approach at the same ages using different promoter fragments allowed the identification of late neurogenic RGCs that can be visualized in the VZ even 24 hours after labeling by electroporation, which produce neurons at later ages. These cells would have been missed in the previous study, given that FlashTag apparently does not label these more quiescent RGCs that remain at early ages for longer times in the VZ. Notably, our data also show that these progenitors have a specific molecular signature with 11 genes being more highly expressed, including Sox9. While we describe here a function for Sox9 for the regulation of differential progenitor behavior, the function of the remaining 10 genes still needs to be investigated.

In summary, we demonstrated here the existence of molecular heterogeneity among neocortical progenitors, which distinguishes two functionally different subpopulations of RGCs. Among the differentially expressed genes that we found, Sox9 has an instructive role in defining neurogenic potential of RGCs. The levels of expression of this gene are further associated with the production of distinct cells types, with higher levels of Sox9 instructing the production of upper layer neurons. These results suggest that the delayed neurogenic behavior instructed by Sox9 is important for preventing some RGCs from early exhaustion, allowing them to generate upper layer neurons at later embryonic ages.

## Acknowledgements

This work was supported by the Spanish Ministry of Science and Innovation (CGS, SAF-2017-82880-R) and NIH (UM, RF1MH121539). C.G.S. holds a Ramón y Cajal Grant from the Spanish Ministry of Science and Innovation (RyC-2015-19058). J.A.M.A. was financed in part by the Coordenação de Aperfeiçoamento de Pessoal de Nível Superior (CAPES, BEX: 717115-3). U.M.is a Bloomberg Distinguished Professor for Neuroscience and Biology. We thank members of the laboratory for comments and criticisms; Isabel Fariñas and Sacramento R. Ferrón for reagents and sharing their equipment with us. We are grateful to Cristina Andrés Carbonell for assistance with mouse maintenance.

## Notes

**Conflict of interest statements:** The authors declare no conflicts of interest

### Competing Interest Statement

The authors have declared no competing interest.

